## Supplementary Information for "Theory and Simulations of condensin mediated loop extrusion in DNA"

Ryota Takaki

*Department of Physics, The university of Texas at Austin*

Atreya Dey, Guang Shi, and D. Thirumalai

*Department of Chemistry, The university of Texas at Austin*

(Dated: August 12, 2021)

**CONTENTS**


I. Simulations 3

A. Model for Condensin 3

B. Energy function 5

C. Simulation with DNA 6

II. Persistence length of condensin 7

III. Derivation of $\boldsymbol{P(L|R)}$ 8

IV. Effect of varying DNA persistence length 10

V. Distribution of LE length per cycle 11

VI. Conformational transition to the LE active state 13

References 15

### I. SIMULATIONS

The purpose of the simulations using a simplified model is to show that the extracted parameter values, obtained by fitting the theoretical extrusion rate or equivalently the velocity of loop extrusion (LE) to experiments, are reasonable. In particular, we use simulations to argue that the value of $\Delta R \approx 26$ nm or equivalently $\sim 76$ bps (see the main text for details) is consistent with experiments that have imaged the shape transitions during the LE process. To this end, we imagine that during the ATPase cycle, the SMC motor undergoes a conformational change from an "open" (top structure in Fig. S1) to a "closed" (bottom structure) state, thus decreasing the distance between the motor domains and the hinge region. We theorize that this process is allosterically driven, in a manner similar to other cargo-carrying motors (myosins, kinesins and dynein), by binding ATP. We envision that in the SMC the allosteric transitions are effectuated through the movement of the flexible elbow region. Such a picture is consistent with high speed AFM imaging [1] and measurements of head-hinge distance as the motor transitions between the two active (open and closed) states [2].

#### A. Model for Condensin

We modeled the two heads of condensin as spheres that are connected by finitely-extensible nonlinear elastic (FENE) potential [3] to the coiled coils (CCs) that connect the motor domains to the hinge (Fig.S1). The CC in the SMCs are reminiscent of the lever arm in Myosin V. The angle $\theta_1$ and $\theta_2$ are formed at the junctions connecting the motor heads to the first bead on the CCs (Fig.S1). As in the well-studied molecular motors, a change in the conformation initiated in the head domain, is amplified to the rest of the motor through the CCs. We envision this process as the principle mechanism by which a spool (roughly $\frac{\Delta R}{0.34} = 76$ base pairs (bps) in a single step) of DNA could be extruded.

We used 19 and 18 beads for upper CC and lower CC, respectively. The diameter of each bead is 1 nm diameter. We used 3 beads (diameter 0.4 nm each) in the middle of the CCs for the elbow region. The strength of the angle potential in the elbow region, marking the break in an otherwise stiff CC, is chosen to facilitate the allosteric propagation of conformational changes in the motor head. For the hinge and two motor heads we used 4 nm diameter

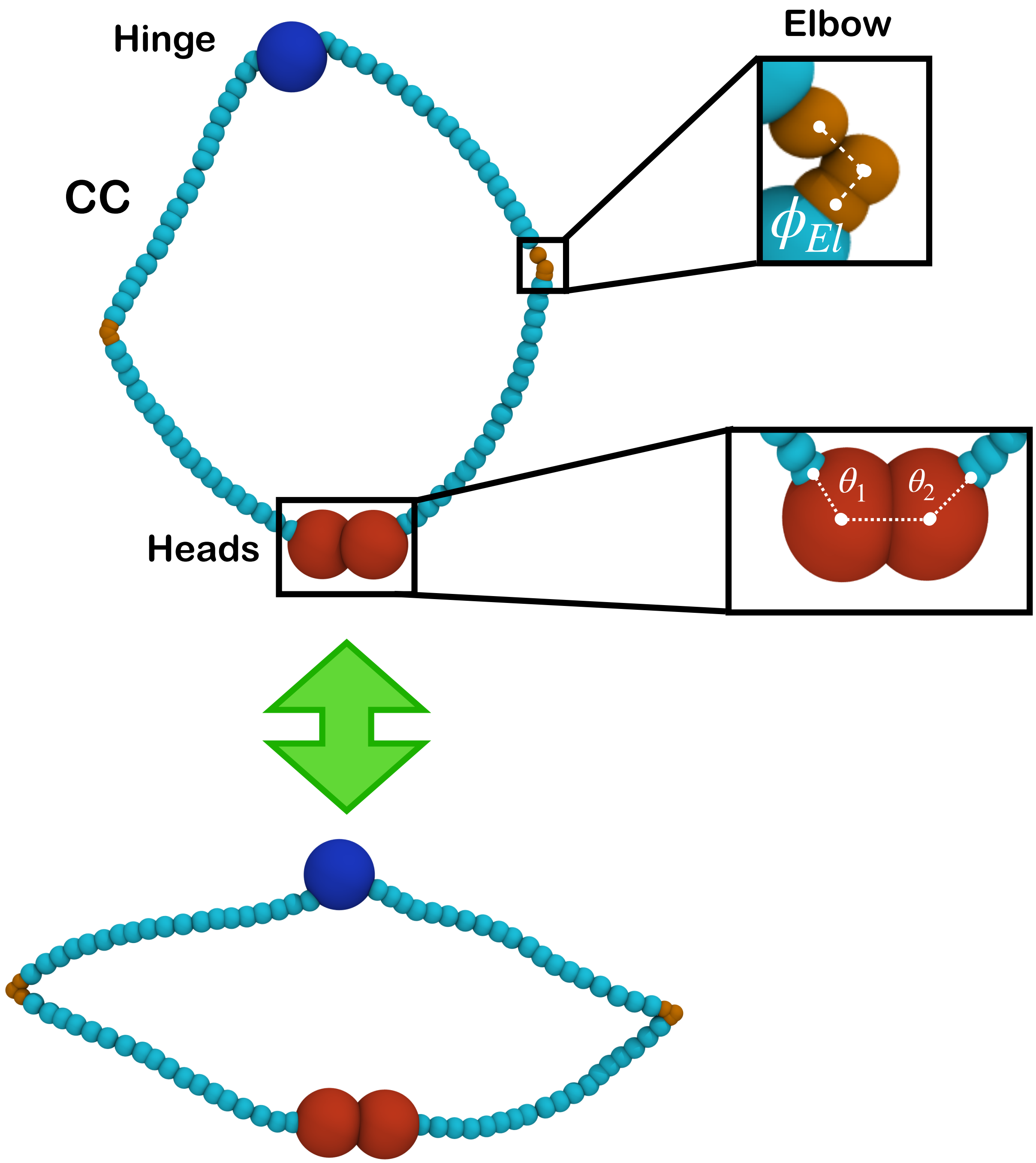


FIG. S1. Cartoon representation of the condensin motor based in part on partially resolved structures and FM images. The simulation model is based on this representation. The two heads are shown as red spheres. A magnified image of the angle between the motor heads at the junctions to the two arms of the SMC is shown in the lower box. The angle at the elbow is depicted in the upper box. The coiled coils (CCs) connecting the motor to the hinge (purple sphere) are treated as a semi-flexible polymers that are kinked at the flexible elbow region. We envision that the allosteric transition between the open and the closed states (shown as by the green arrow) is driven by ATP binding to the motor domains.

beads.

All the lengths are measured in units of $\sigma = 1$ nm, corresponding to the diameter of the beads in the CC. We express energy in $k_BT$ units, where $k_B$ is the Boltzmann constant and $T$ is the temperature. The mass of all the particles were set to $m = 1$. We performed low-friction Langevin dynamics simulations using OpenMM [4] software using a time-step of $\Delta t_L = 0.01\tau_L$, where $\tau_L = 0.4\sqrt{m\sigma^2/k_BT}$. The value of the friction coefficient is $0.01/\tau_L$.

#### B. Energy function

Because our goal is to merely illustrate that the proposed allosteric mechanism for SMC-mediated LE is plausible, we chose a simple energy function to monitor the conformational changes in condensin. The explicit form of the energy function is,

$$E(\vec{r}_1, \vec{r}_2, ..., \vec{r}_N, \vec{\phi}, \vec{\theta}) = \sum_{i=1}^{N-1} U_{FENE}(r_{i,i+1}) + \sum_{i\neq j}^{N} U_N(r_{i,j}) + \Big( \sum_{i\in CC\neq El} U_{ANG}^{CC}(\phi_i) + \sum_{i\in El} U_{ANG}^{El}(\phi_i) \Big) + \sum_{i\in Head} U_{CNF}(\theta_i). \tag{S1}$$

The first term in Eq.(S1) enforces the connectivity of the beads and is given by,

$$U_{FENE}(r_{i,i+1}) = -\frac{1}{2}k_F R_F^2 \log\Big[1 - \frac{(r_{i,i+1} - r_{i,i+1}^0)^2}{R_F^2}\Big], \tag{S2}$$

where $k_F$ is the stiffness of the potential, $R_F$ is the upper bound for the displacement, and $r_{i,i+1}^0$ is the equilibrium distance between the beads, $i$ and $i+1$. The second term in Eq.(S1), accounting for excluded volume interactions, is given by,

$$U_N(r_{i,j}) = \epsilon_N \Big(\frac{\sigma}{r_{i,j}}\Big)^{12}, \tag{S3}$$

where $\epsilon_N$ and $\sigma$ are the strength and range of the interaction, respectively. We used additive interactions, which means that $\sigma$ is the sum of the radii of the two interacting beads. The third and the fourth terms in Eq.(S1) are the two angle potentials that control the bending stiffness of the CCs. The potential, $U_{ANG}(\phi_i)$, is taken to be,

$$U_{ANG}(\phi_i) = \epsilon_b(1 + \cos\phi_i), \tag{S4}$$

where $\epsilon_b$, the energy scale for bending, is related to the persistence length of the semi-flexible CC. We used a smaller value of $\epsilon_b$ for $U^{El}_{ANG}(\epsilon^{El}_b)$ to model the difference in the stiffness between the elbow region, and the rest of the CC.

The last term in Eq.(S1) models the conformation change in the motor head due to ATP binding, and is taken as,

$$U_{CNF}(\theta_i) = \frac{1}{2}k_C\left(\theta^0_i - \theta_i\right)^2, \tag{S5}$$

where $k_C$ is the spring constant for the potential, and $\theta^0_i$ is the equilibrium angle for the angle potential. Before the conformational change, we set $\theta^0_i = 2.4$ (radian) in the open state, which is roughly the angle calculated by ATP engaged state of prokaryotic SMC [5]. Because the structure for the closed state is unavailable, we chose $\theta^0_i = 4.0$ (radian) for the closed state, which leads to $\theta_i \sim \pi$ (radian) in equilibrium. The transition between the open and closed states results in the scrunching of the DNA (the nearly stationary motor reels in the DNA), and extrusion of the loop. The parameter values in the energy function used in the simulations are in Table S1.

##### C. Simulation with DNA

We used a coarse-grained bead-spring model for DNA [6, 7]. Each bead represents 10 base-pairs, which implies that the bead size is $\sigma_{DNA} = 3.4$ nm. The chain has $N = 100$ beads or 1,000 base-pairs. Consecutive beads along the chain are connected by a FENE potential (Eq.(S2)) with $r^0_{i,i+1} = \sigma_{DNA}$. The DNA stiffness is modeled using a harmonic bending potential given by,

$$U_{BEND} = \frac{1}{2}\sum_{i=1}^{N_{ang}} \alpha \cdot \theta^2, \tag{S6}$$

where $N_{ang} = 98$ is the number of bond angles, $\theta_i$ is the deviation of the $i^{th}$ bond-angle in the chain from 180 deg, and $\alpha = 15.55$ ($k_BT/\text{rad}^2$) is a constant [6]. We chose $\alpha$ so that the persistence length is $\approx 50$ nm, which is the canonical value for DNA in monovalent salts.

| Parameter | Value |
| --- | --- |
| $k_F$ | $50(k_BT/\mathrm{nm}^2)$ |
| $R_F$ | $1.5(\mathrm{nm})$ |
| $\epsilon_b^{CC}$ | $150(k_BT)$ |
| $\epsilon_b^{El}$ | $2(k_BT)^*$ |
| $\epsilon_N$ | $5(k_BT)$ |
| $k_C$ | $200(k_BT/\mathrm{rad}^2)$ |

TABLE S1. Parameters for the molecular dynamics simulation. * The dependence of the distance distribution between the head and hinge during one cycle on $\epsilon_b^{El}$ is given in Fig. S3.

#### II. PERSISTENCE LENGTH OF CONDENSIN

In order for the allosteric transition that brings the head-hinge distance to within $\Delta R \sim 26$ nm ($\approx 22$ nm in experiments and simulations), condensin has to be sufficiently rigid but not overly so. We estimate the persistence length of condensin ($l_p^{CC}$) by fitting the theoretical expression for end-to-end distance ($R$) with contour length $L$ [8] to the simulation results. Here, $R$ is the head-hinge distance in the O shape and $L = 51$ nm is the length of the CC in the simulation. As we varied $\epsilon_b^{El}$, $l_p^{CC}$ changes, resulting in different $\Delta R_s$ values (Fig. S3). For $l_p^{CC} \approx 24$ nm, we find that $\Delta R_s = 22$ nm, which coincides with the experimental value [2]. The value of $l_p^{CC}$ is roughly six times larger than the experimental data. This may be due to the differences in the fitting used in the simulations or the errors in accurate measurements or in the method used to extract $l_p^{CC}$ from the experimental data. It should be emphasized that for comparison between theory and experiment $l_p^{CC}$ does not play an important rule. Nevertheless, it would be most interesting to design experiments to obtain precise estimates of $l_p^{CC}$.

The results in the inset in Fig.S3(a) are calculated using $l_p^{CC} \sim 4$ nm. In this case, the elbow effect is eliminated by setting $\epsilon_b^{CC} = \epsilon_b^{El} = 4k_BT$, making the entire CCs flexible. The simulations show that $\Delta R \approx 15$ nm, which disagrees with the experimental value ($\Delta R \approx 22$ nm). We infer that the existence of rigid portion of CC with flexibility in the elbow region is essential for the scrunching mechanism.

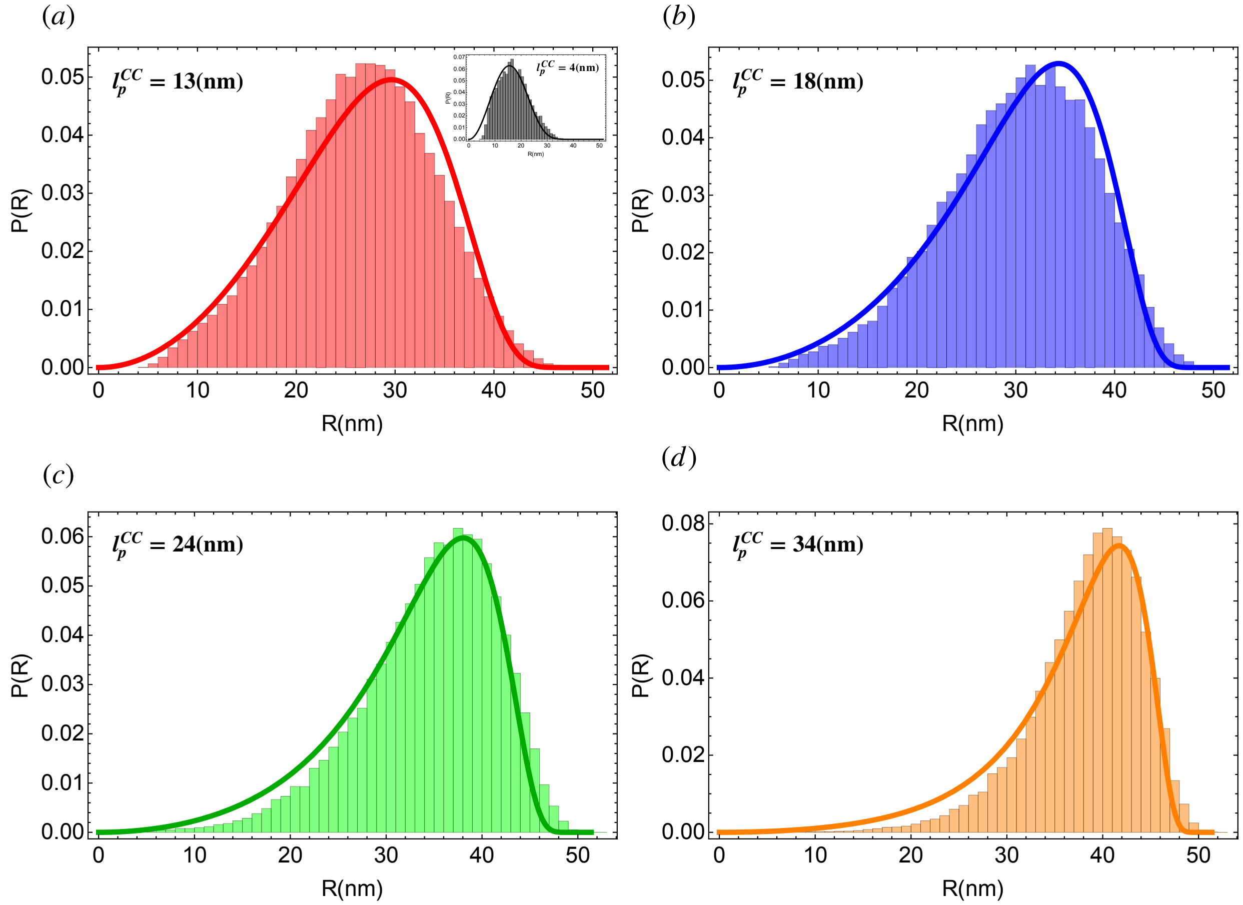


FIG. S2. Extracting $l_p^{CC}$ from the simulations. Histograms are for the head-hinge distance in the O shape from the simulations. Solid lines are from the theoretical expression for $P(R)$. The value of $\epsilon_b^{CC} = 150(k_BT)$ in all the panels except in the inset of (a). (a) $l_p^{CC} \sim 13$ nm for $\epsilon_b^{El} = 0(k_BT)$. Inset: $l_p^{CC} \sim 4$ nm for $\epsilon_b^{CC} = 4(k_BT)$ and $\epsilon_b^{El} = 4(k_BT)$. (b) $l_p^{CC} \sim 18$ nm and $\epsilon_b^{El} = 1(k_BT)$. (c) $l_p^{CC} \sim 24$ nm and $\epsilon_b^{El} = 2(k_BT)$. (d) $l_p^{CC} \sim 34$ nm and $\epsilon_b^{El} = 3(k_BT)$.

#### III. DERIVATION OF $P(L|R)$

A major ingredient in the theory (see Eq.(1) in the main text) is the calculation of the contour length of the extruded loop as condensin is powered by ATP binding to the motor head, followed by hydrolysis, and subsequently resetting to complete the catalytic cycle. To obtain Eq.(1) in the main text, let us consider condensin separated by the spatial distance $r$ that pinches a loop whose genomic length is $s$ (Fig.S4). Given the distribution of the spatial distance $r$ between two loci separated by a linear genomic distance $s$, $P(r|s)$, we would like to derive the distribution of $s$, $P(s|r)$. Indeed, $P(s|r)$ is the probability density of the extruded length of DNA, $s$, by condensin whose DNA binding domains are separated

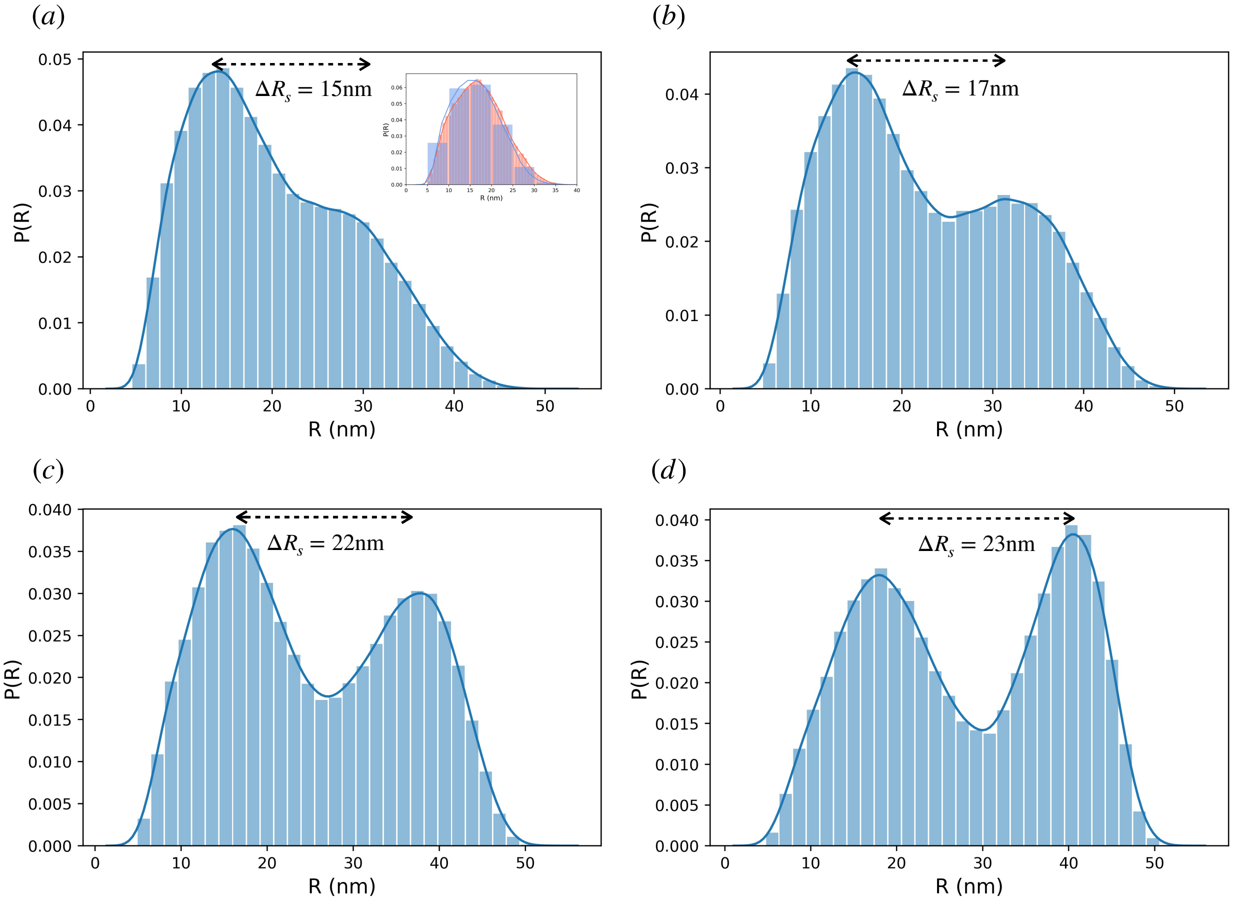


FIG. S3. $P(R)$ for different values of $l_p^{CC}$ in the simulations. (a) $l_p^{CC} \sim 13$ nm $[\epsilon_b^{El} = 0(k_BT)]$. The position of the peaks are $R_2 \sim 13$ nm and $R_1 \sim 28$ nm. Inset: $l_p^{CC} \sim 4$ nm $[\epsilon_b^{CC} = \epsilon_b^{El} = 4(k_BT)]$. The position of the peaks are $R_2 \sim 15$ nm and $R_1 \sim 16$ nm. (b) $l_p^{CC} \sim 18$ nm $[\epsilon_b^{El} = 1(k_BT)]$. The position of the peaks are $R_2 \sim 14$ nm and $R_1 \sim 32$ nm. (c) $l_p^{CC} \sim 24$ nm $[\epsilon_b^{El} = 2(k_BT)]$. The position of the peaks are $R_2 \sim 16$ nm and $R_1 \sim 38$ nm. (d) $l_p^{CC} \sim 34$ nm $[\epsilon_b^{El} = 4(k_BT)]$. The peak positions are $R_2 \sim 18$ nm and $R_1 \sim 40$ nm.

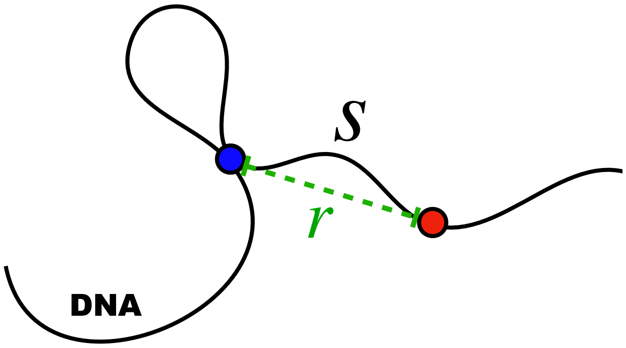


FIG. S4. A picture of a conformation of condensin bound to two loci separated by a genomic distance $s$ (extruded loop length). The spatial distance between the attachment points in the DNA is $r$. For LE to occur condensin has to engage with at least two loci on the DNA.

by the distance, $r$. According to the Bayes theorem, we have $P(r|s)P(s) = P(s|r)P(r)$. The normalization dictates that $P(r) = \int_0^L P(r|s)P(s)\mathrm{d}s$. These two equations lead to,

$$P(s|r) = \frac{P(r|s)P(s)}{\int_0^L P(r|s)P(s)\mathrm{d}s}.$$

We assume that there is no preference for picking a specific genomic distance $s$ on the DNA to which the motor attaches to initiate the extrusion process. Consequently, we take $P(s) = 1/L$. Therefore,

$$\begin{aligned} P(s|r) &= \frac{(1/L)P(r|s)}{(1/L)\int_0^L P(r|s)\mathrm{d}s} \\ &= \frac{P(r|s)}{\int_0^L P(r|s)\mathrm{d}s}. \end{aligned}$$

Thus, $P(s|r)$ and $P(r|s)$ differ only by a constant, $\int_0^L P(r|s)\mathrm{d}s$, if we consider a fixed $r$. It is clear that $P(r|s)$ is the radial probability density for the interior segments separated by a distance $r$ for a semi-flexible polymer, which is derived elsewhere [6]. For the case $s = L$, $P(r|s)$ is the result for the distribution of end-to-end distance for semi-flexible chains, $P(R|L)$ [8]. It is known that the simple analytic result for $P(R|L)$ [8] is accurate when compared to the exact result [9] or numerical simulations. Thus, we employ the simpler expression $P(R|L)$ and assume that $P(L|R)$ is equivalent to $P(R|L)$ up to a normalization constant when expressed in terms of $L$ with fixed $R$. Calculation of the distribution of loop sizes requires knowing, $P(L|R, f)$, which can also be derived from $P(R, f|L)$ in a similar manner.

#### IV. EFFECT OF VARYING DNA PERSISTENCE LENGTH

In the main text, we used $l_p = 50$ nm (∼147 bps) as the persistence length of DNA, which is widely accepted value for DNA [10]. It could be interesting to explore the consequences of varying $l_p$, which can be drastically altered in the presence of divalent cations, as a variable in our theory. In Fig.S5(a) we plotted $P(L|R = 50\text{ nm})$ using Eq.(1) in the main text for different $l_p$. As DNA becomes flexible the distribution of $P(L|R = 50\text{ nm})$ becomes wider, suggesting that most probable value of the captured length of DNA by condensin would be

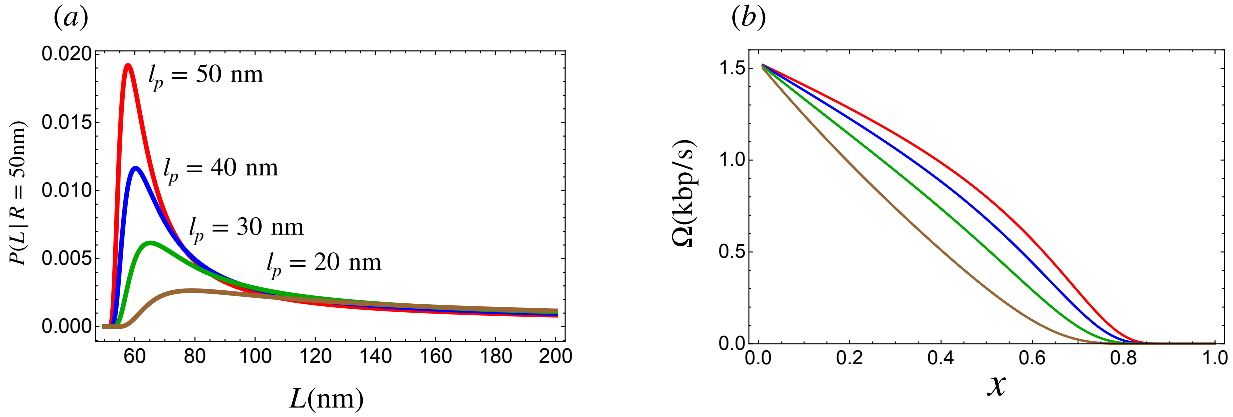


FIG. S5. Effect of variable persistence length for DNA. (a) Plot of $P(L|R = 50$ nm$)$. $l_p = 50$ nm (red), $l_p = 40$ nm (blue), $l_p = 30$ nm (green), and $l_p = 20$ nm (brown). (b) Extrusion rate of DNA for different $l_p$. $l_p = 50$ nm (red), $l_p = 40$ nm (blue), $l_p = 30$ nm (green), and $l_p = 20$ nm (brown). $\Delta R$ is fixed to be 26 nm $\sim$ 76 bps.

larger with a large dispersion. Thus, in this situation our approximation, $L \approx R$, would become less accurate. Nevertheless, we can explore the velocity of extrusion for different $l_p$ shown in Fig.S5(b) for a fixed $\Delta R = 26$ nm. As $l_p$ decreases, the velocity of extrusion becomes linear and slower because the load acting on DNA is higher for smaller $l_p$ at the same extension.The decrease of $\Omega$ as $l_p$ decreases can be deduced from the linear (small $x$) expansion of Eq.(7) in the main text. At small forces we find that $f = \frac{3k_BT}{2l_p}$. Substituting this linear expansion in the expression for $\Omega$ [Eq.(6)] confirms that as $l_p$ decreases $\Omega$ becomes smaller. Note that this holds for fixed extrusion length per step ($\sim$26 nm obtained by fitting Eq. (6) in the main text to the measured LE velocity).

#### V. DISTRIBUTION OF LE LENGTH PER CYCLE

In the main text, we showed that the theoretically derived distribution of LE length agrees well with the one obtained in the experiment [11] for $f = 0.4$ pN, *without adjusting any parameters*. The agreements persist for different values of load unless $0.5$ pN $\leq f$ (Fig.S6). The discrepancies for high loads can be eased by using smaller $\Delta$ as shown in the blue distributions. It is worth mentioning that the sample sizes for obtaining the experimental results in Fig. S6 are much smaller than for $f = 0.4$ pN. They are $N = 153, 131, 102, 118, 155, 140$ for $f = 0.2$ pN$, 0.3$ pN$, 0.5$ pN$, 0.6$ pN$, 0.7$ pN$, 1.0$ pN respectively whereas $N = 1727$ for $f = 0.4$ pN in the main text. The smaller sample sizes surely affects the accuracy of the

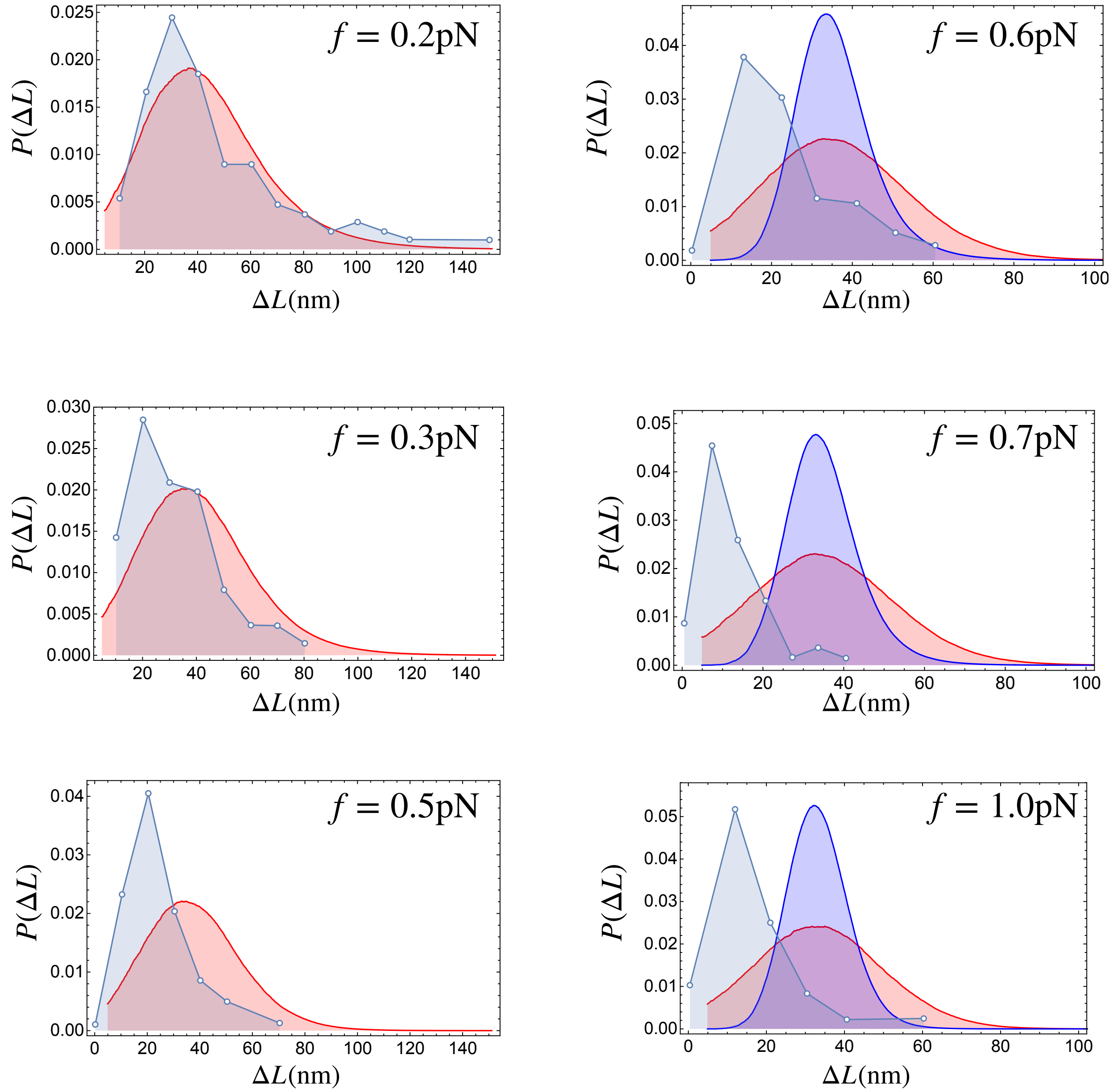


FIG. S6. Distributions for LE length per step for various external loads on DNA. The points in blue are from the experiment [11] and the distributions in red are from the theory (main text). The distributions in blue for high external loads are for the different standard deviation ($\Delta = 5$ nm) for $R_1$.

measured distribution. Considering the low stall load for condensin, likely to be $f \approx 0.8$ pN (see the inset of Fig.4a in the main text), we believe that our theory predicts, with reasonable accuracy, the LE length by condensin during DNA compaction.

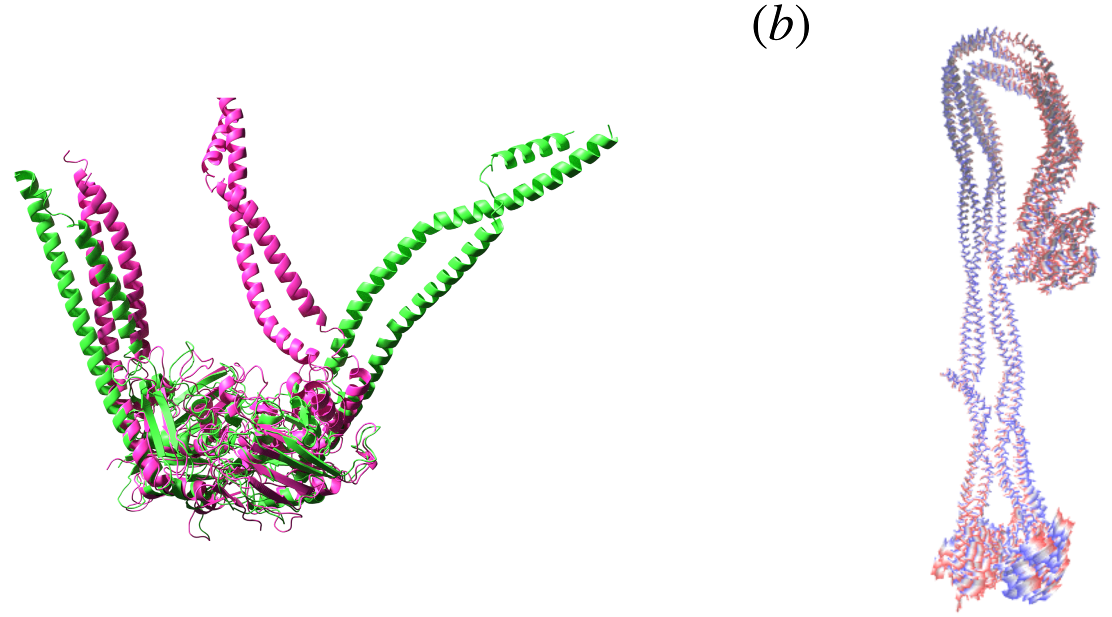


FIG. S7. (a) Structural alignment obtained by minimizing the root mean squared displacement for the head domain. Structure in green is the *apo* state (PDB:6YVU) where the heads are aligned to the ATP bound state, and the structure in magenta is the state with ATP (PDB:6YVD). Alignment is constructed only for the heads. (b) Normal mode analysis using the crystal structure for the *apo* state and ATP bound state (6YVU and 6YVD, respectively). Blurred regime shows the possible movements of the residues.

#### VI. CONFORMATIONAL TRANSITION TO THE LE ACTIVE STATE

The lack of structures in various nucleotide states of condensin in the presence of DNA makes it virtually impossible to provide a molecular basis of loop extrusion. In the absence of viable structures, and prompted by the theoretical prediction that during a single turnover the distance between the hinge and motor should come within $\Delta R \approx (22 - 26)$ nm, we envisioned that the SMCs transition between the open and closed states in order to extrude loops. This picture is fully consistent with experiments [2]. The shape of condensin from the partial structures of condensin in the inactive (in the absence of nucleotides or DNA) obtained by cryo-EM at $\sim 8.1$Å resolution [12] cannot account for the O shape, which likely represents the functionally active state. Because the cryo-EM structures [12] do not contain DNA, they correspond to an inactive state. Nevertheless, even these inactive structures [12] reveal that binding of ATP, sandwiched between the two heads, leads to a substantial conformational change ($\sim$ 50 rotation and $\sim$ 20Å translation) near the junction between the CC and the heads.

To provide insights into the ATP-induced conformational changes, we first aligned the structures in the head-domains of the *apo* state (PDB:6YVU) and the head-domains of the ATP-bound state (PDB:6YVD). Using VMD's multiseq tool [13], we aligned residues: 1-148, 1036-1170 for Chain A and 150-311, 1285-1415 for Chain B using STAMP structural alignment. The structural alignment shows that there is a large change in the CC orientation between the *apo* state and the ATP bound state. Fig. S7a shows the structural alignment obtained by minimizing the root mean squared displacement between the head domains in the *apo* state (PDB:6YVU), and the head domain in the ATP bound state (PDB:6YVD). The alignment suggests that the CC could undergo a wide opening motion upon ATP binding.

**Normal mode analysis:** To understand the correlation between the head and hinge movement, we performed Normal Mode Analysis on the condensin structure in the *apo* state (PDB:6YVU). We created a variant of an Elastic Network Model using a second-order Taylor series expansion of the Self-organized Polymer model with Side-chains (SOP-SC) for proteins [14]. Since the *apo*-aligned to ATP bound state does not have the full condensin structure, we calculated a displacement vector ($\mathbf{D}$) between 6YVU (aligned to 6YVD) with the original coordinate of 6YVU using only the beads that were present in both the full condensin structure, and the *apo*-aligned structure. Using the normal modes, we determined the eigenvector that best approximates this structural transition. $\mathbf{D}$ is a $3N$ dimensional vector where N is the total number of beads (PDB: 6YVD). $\mathbf{D}_i = [(\mathbf{D})_{1,x}, ..., (\mathbf{D})_{M,z}]$, where $(\mathbf{D})_{j,\alpha}$ is the entry associated with bead j, and direction $\alpha \in x, y, z$. The beads which are not present in the ATP-aligned structure have $(\mathbf{D})_{j,\alpha} = 0$. The overlap between the eigenvector $v_n$ (corresponding to normal mode $n$), and displacement $\mathbf{D}$ is given by,

$$I_n(\mathbf{D}) = \frac{\sum_{i=1}^{3N} v_{n,i} \cdot D_i}{\sqrt{\sum_{i=1}^{3N} v_{n,i}^2}\sqrt{\sum_{i=1}^{3N} \mathbf{D}_i^2}}. \tag{S7}$$

We found that mode 7 has the highest overlap value.

Fig. S7b shows the motion of the residues calculated from the largest eigenvector that produces the maximum overlap. In the blurred region residues experience correlated movement. Our analysis shows that the normal modes mostly overlap at the head and hinge. In other words, head and hinge are likely to undergo the largest conformational change during the transition between the two states. If this finding holds when the structures of condensin in various nucleotide binding states are determined, it would provide a molecular basis for the O and B shapes that the condensin clearly samples during the process of loop extrusion [2].

---

[14] M. L. Mugnai, C. Templeton, R. Elber, and D. Thirumalai, bioRxiv (2020).
